# Discovery of novel haplotypes for complex traits in landraces

**DOI:** 10.1101/2020.05.25.114264

**Authors:** Manfred Mayer, Armin C. Hölker, Eric González-Segovia, Thomas Presterl, Milena Ouzunova, Albrecht E. Melchinger, Chris-Carolin Schön

## Abstract

Genetic variation is of crucial importance for selection and genetic improvement of crops. Landraces are valuable sources of diversity for germplasm improvement, but for quantitative traits efficient strategies for their targeted utilization are lacking. Here, we propose a genome-based strategy for making native diversity accessible for traits with limited genetic variation in elite germplasm. We generated ~ 1,000 doubled-haploid (DH) lines from three European maize landraces, pre-selected based on molecular and phenotypic information. Using GWAS, we mapped haplotype-trait associations for early development traits at high resolution in eleven environments. Molecular haplotype inventories of landrace derived DH libraries and a broad panel of 65 breeding lines based on 501,124 SNPs revealed novel variation for target traits in the landraces. DH lines carrying these novel haplotypes outperformed breeding lines not carrying the respective haplotypes. Most haplotypes associated with target traits showed stable effects across populations and environments and only limited correlated effects with undesired traits making them ideal for introgression into elite germplasm. Our strategy was successful in linking molecular variation to meaningful phenotypes and identifying novel variation for quantitative traits in plant genetic resources.

## Introduction

Harnessing the allelic diversity of genetic resources is considered essential for overcoming the challenges of climate change and for meeting future demands on crop production^1,2^. For most traits of agronomic importance, modern breeding material captures only a fraction of the available diversity within crop species^1^. In the case of maize (*Zea mays* L.), today’s elite germplasm went through several bottlenecks, first by geographical dispersion from its center of origin^3,4^, second through the selection of only a few key ancestors sampled from a small number of landraces to establish heterotic groups^5,6^, and third through decades of advanced cycle breeding with high selection intensities^7,8^. For traits that were not targets of selection in the past but are important today, like abiotic stress tolerance and resource-use efficiency^9^, this might have resulted in the loss of favorable alleles during the breeding process. In addition, unfavorable alleles might have become fixed during the selection process due to drift and/or hitchhiking effects^10–12^.

Impressive examples exist where introgression of novel alleles from genetic resources have improved mono- or oligogenic traits^13–15^, but for broadening the genetic diversity of complex traits such as yield or abiotic stress tolerance successful examples are scarce^2^. Up to date, the genomic characterization of genetic resources was predominantly based on sampling individuals across a wide range of accessions, maximizing the level of diversity in the genetic material under study^2,16–20^. Such diverse samples are characterized by high variation in adaptive traits and strong population structure, leading to spurious associations and limited power for detecting associations with non-adaptive traits of agronomic importance^21,22^. Furthermore, novel alleles which are locally common but globally rare likely remain undetected in broad, species-wide samples, whereas in a more targeted approach they might show sufficiently high frequencies for detection^22^.

Here, we propose a genome-based strategy (Supplementary Fig. 1) for making native diversity of maize landraces accessible for improving quantitative traits showing limited genetic variation in elite germplasm, such as cold tolerance and early plant development^23–25^. Capitalizing on low levels of linkage disequilibrium (LD) we mapped haplotype-trait associations at high resolution in ~ 1,000 doubled-haploid (DH) lines derived from three European flint maize landraces. The genetic material was pre-selected for adaptation to target environments to avoid confounding effects of strong adaptive alleles as suggested by Mayer et al.^26^. Novelty of promising haplotypes was assessed genotypically by quantifying their frequency in a diverse panel of 65 European flint breeding lines. Phenotypically, the direction and magnitude of haplotype effects was evaluated relative to a subset of breeding lines. Many of the discovered novel haplotypes showed stable trait associations across populations and environments. In addition, for most of them no undesired trait associations were observed, making them ideal for introgression into elite germplasm. We show that our strategy to sample comprehensively individuals from a limited set of pre-selected landraces was successful in linking molecular variation to meaningful phenotypes and in identifying novel alleles for quantitative traits in plant genetic resources.

## Results

### Novel variation from maize landraces

Our goal was to investigate if three DH libraries derived from the pre-selected landraces Kemater Landmais Gelb (KE), Lalin (LL) and Petkuser Ferdinand Rot (PE) carried novel alleles compared to a diverse panel of 65 European breeding lines, representing a large number of different source populations across Europe^27^. We first performed a principal coordinate analysis (PCoA) based on 501,124 single nucleotide polymorphism (SNP) markers jointly for the full set of 941 landrace derived DH and 65 European breeding lines (Fig. 1a). The first principal coordinate explained 6.2% of the molecular variation and separated the landrace derived and the breeding lines based on their geographical origin within Europe from north-east (Germany) to south-west (southern France, Spain). The second principal coordinate explained 5.4% of the variation and separated the two landraces PE and KE from the panel of breeding lines.

**Fig. 1:**
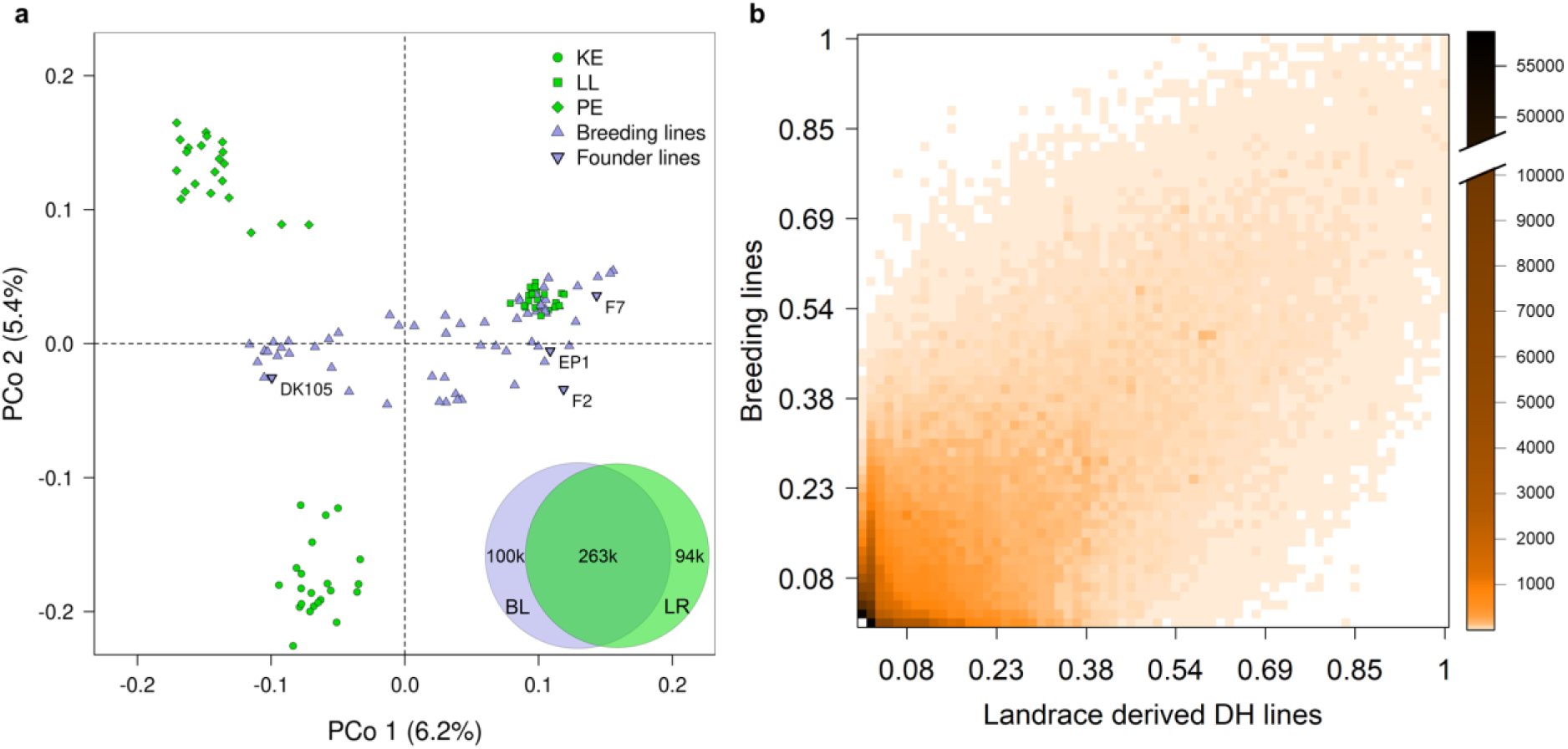
Molecular inventories point to novel variation in landraces. **(a)** Principal coordinate analysis based on pairwise Modified Rogers’ distances of 66 landrace derived DH lines and 65 breeding lines. From each of three DH libraries (KE, LL and PE) 22 lines were sampled randomly. Axis labels show the percentage variance explained per principal coordinate. Venn diagram shows overlap of 456,911 haplotypes between 941 landrace derived DH lines (LR) and 65 European breeding lines (BL). Haplotypes were constructed for non-overlapping genomic windows of 10 SNPs. **(b)** Frequency of 456,911 haplotypes in DH lines (x-axis) and breeding lines (y-axis). Colors indicate the number of haplotypes within each cell of the heat map.

In addition to the PCoA we constructed haplotypes in non-overlapping genomic windows of ten SNPs for the 941 landrace derived DH lines and the 65 European breeding lines. In total, the landrace and breeding line panels comprised 356,724 and 363,290 haplotypes (Fig. 1a) corresponding to an average of 7.12 and 7.25 haplotypes per window, respectively. As expected for genetic material originating from the same germplasm group (European flint maize), haplotype frequencies were positively correlated (Pearson’s *r* = 0.74, P < 2.2e-16) between the two panels (Fig. 1b). Overall, 26.2% of the haplotypes of the landrace panel were not present in the breeding lines, indicating novel haplotype variation. For those haplotypes median and mean frequencies in the landrace panel were 0.005 and 0.039, respectively. Only 2.7% of those haplotypes occurred in all three landraces, whereas 82.8% occurred in only one landrace. Within the respective individual landraces their median and mean frequencies increased to 0.065 and 0.101, respectively. The landrace panel captured 72.4% of the haplotypes present in the panel of breeding lines.

### Trait-associated genomic regions

To evaluate if molecular inventories of landrace derived material are predictive for their potential to improve traits of agronomic importance, we performed haplotype based genome-wide association scans (GWAS) for nine traits. Trait-associated genomic regions were defined based on LD between significant haplotypes (Methods; Fig. 2, Supplementary Table 1). As landraces were pre-selected for variation in early plant development^26,28^, most associations (37 to 55) were detected for the traits early vigor (EV_V4/V6) and early plant height (PH_V4/V6).

**Fig. 2:**
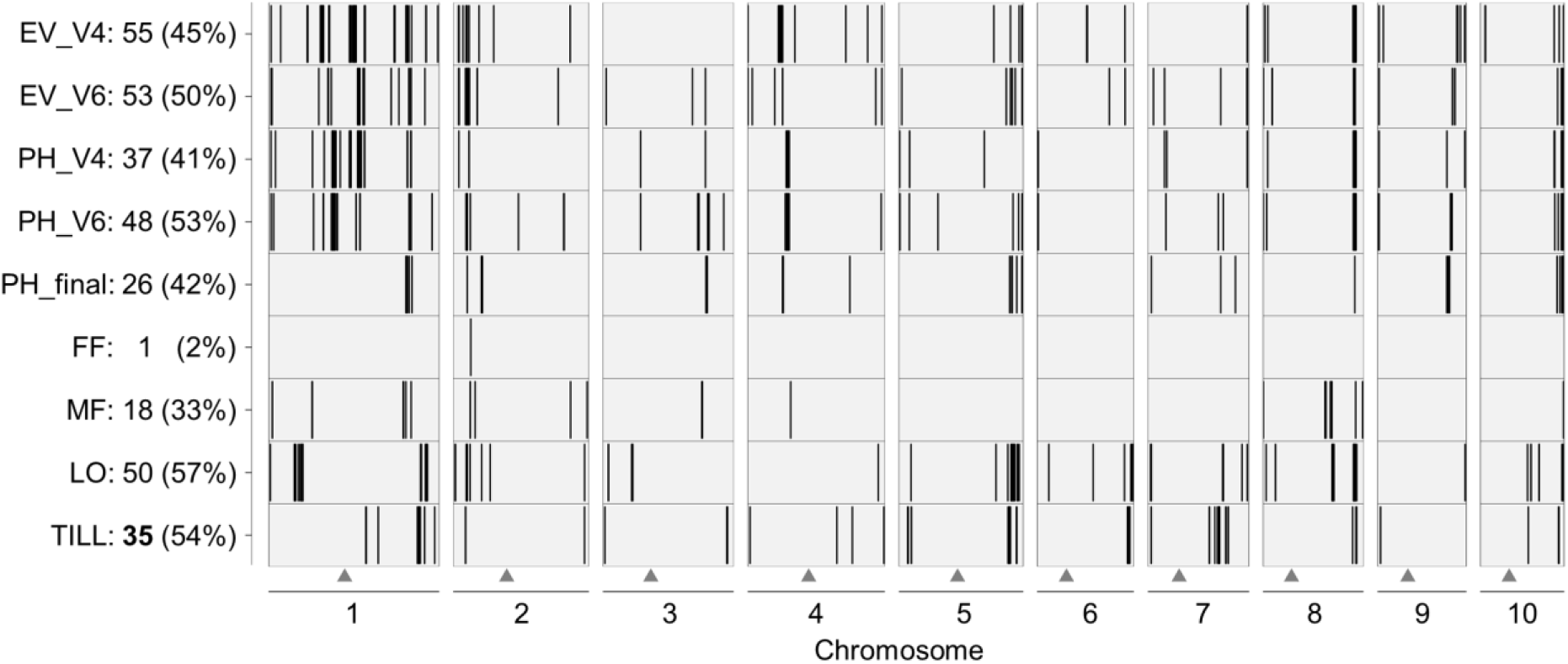
Results from GWAS in DH-libraries derived from maize landraces. Black vertical bars indicate the position of genomic regions significantly associated with nine traits (y-axis) in 899 landrace derived DH lines. The x-axis shows the ten chromosomes of maize. Triangles mark the position of the centromere for each chromosome. The y-axis indicates the trait, the number of significant regions per trait, and the percentage genetic variance explained.

Haplotypes explained between 2% (female flowering time, FF) and 57% (lodging, LO) of the total genetic variance of the respective traits (Fig. 2). Despite the large sample size (n = 899), the proportion of genetic variance explained might be somewhat overestimated^29,30^ and thus has to be interpreted with caution. Only few genomic regions were detected for flowering time indicating that alleles with large effects were fixed during adaptation of the respective landraces to their geographical region, thus having little impact on GWAS for other traits.

Average *r*^*2*^ decay distances (*r*^*2*^ < 0.2) within the three DH libraries were 203 (LL), 484 (PE) and 973 kb (KE), and 201 kb for the combined set. This is consistent with previous results^26^ and warrants high mapping resolution in the three DH libraries under study. For comparison, the diverse panel of 65 breeding lines across Europe exhibited an average *r*^*2*^ decay distance of 107 kb. The lower LD level in the breeding line panel can be explained by admixture of many different source populations with varying linkage phases, which is generally undesired in GWAS. The median size of genomic regions associated with the nine traits under study was 92 kb, with a median number of three annotated genes per region (Supplementary Fig. 2), enabling prediction of candidate genes and functional analyses. Only for a few regions (< 5%) resolution was not optimal as they comprised more than 100 annotated genes. Mapping resolution in the three DH libraries is best demonstrated by an example of an already well characterized locus: *teosinte branched 1* (*tb1*). The gene *tb1* played a major role in the transition from highly branched teosinte to maize with strongly reduced branch development^31^. In our study, a strong significant association for tillering (TILL) was found in a genomic region comprising the *tb1* locus (size 1.3 Mb, including in total 22 genes; Supplementary Table 1). *In silico* fine mapping in the respective region (Methods) identified a 10 SNP window which overlapped perfectly with *tb1* and its regulatory upstream region.

### Effect size and stability of trait-associated haplotypes

The potential of the identified haplotypes for elite germplasm improvement depends on the size and direction of their effects on the traits of interest, their environmental stability and their dependence on the genetic background. In a given trait-associated genomic region one window of 10 SNPs comprising several haplotypes was selected. Significant haplotypes, hereafter referred to as focus haplotypes, entered into a multi-environment model (Supplementary Fig. 3) and were classified into “favorable”, “unfavorable” and “interacting” based on the direction and stability of their effects in the different test environments (Supplementary Fig. 4). According to this categorization scheme, a high number of favorable haplotypes for early plant development traits were found in the DH libraries (Table 1, Fig. 3a), representing potential candidates for introgression into elite germplasm. For the undesirable traits LO and TILL, most identified haplotypes were unfavorable. Overall, haplotypes identified for all nine traits showed moderate to high effect stability across environments, with similar patterns for favorable and unfavorable haplotypes (Fig. 3a,b).

**Table 1:** Number and percentage of favorable, unfavorable and interacting focus haplotypes per trait. Haplotypes with consistent effect direction across environments were categorized as favorable or unfavorable. For EV_V4, EV_V6, PH_V4 and PH_V6 positive (negative) effects were defined as favorable (unfavorable). For LO and TILL negative (positive) effects were defined as favorable (unfavorable). Haplotypes with changing effect direction were categorized as interacting.

| Trait | Equal effect direction across environments |  | Changing effect direction across environments |
| --- | --- | --- | --- |
|  | Favorable, N (%) | Unfavorable, N (%) | Interacting, N (%) |
| EV_V4 | 16 (29%) | 29 (53%) | 10 (18%) |
| EV_V6 | 14 (26%) | 26 (49%) | 13 (25%) |
| PH_V4 | 15 (41%) | 15 (41%) | 7 (19%) |
| PH_V6 | 20 (42%) | 22 (46%) | 6 (13%) |
| LO | 11 (22%) | 35 (70%) | 4 (8%) |
| TILL | 11 (31%) | 23 (66%) | 1 (3%) |

**Fig. 3:**
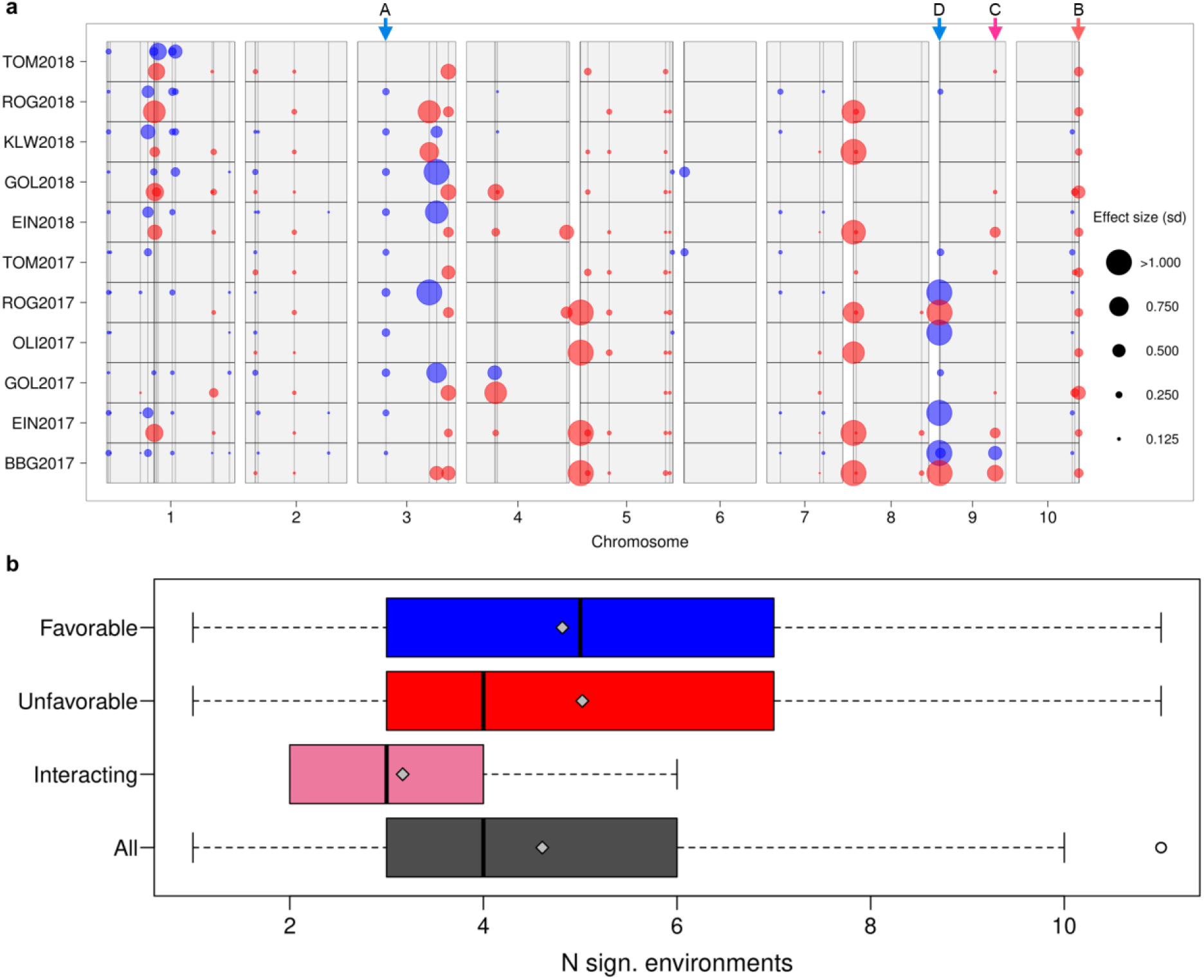
Effect stability of focus haplotypes across environments. **(a)** Genomic position as well as effect size and direction for 48 haplotypes associated with PH_V6 across 11 environments. Circles indicate significant haplotypes with effect sizes given in phenotypic standard deviations. Positive and negative effects are colored in blue and red, respectively. Arrows at the top indicate the positions of haplotypes described in Supplementary Fig. 4. **(b)** Number of environments in which favorable (n = 65), unfavorable (n = 93), interacting (n = 36) and all (n = 194) haplotypes had significant effects on four early plant development traits (EV_V4, EV_V6, PH_V4 and/or PH_V6). Gray diamonds indicate means.

To evaluate the dependency of haplotype effects on the genomic background, we compared effect significance and sign of the identified focus haplotypes between landraces KE and PE. From the 48 haplotypes associated with PH_V6, comparisons could be made for 19 haplotypes present in both KE and PE. Together, these 19 haplotypes showed 115 environment-specific haplotype-trait associations, from which 35 (30%) were significant for both landraces (Supplementary Fig. 5a). All of those 35 associations had equal effect signs for both landraces. Also for the 80 environment-specific associations significant for only one of the two landraces, a large majority (90%) had equal effect signs for both landraces. Similar patterns were observed for PH_V4 (Supplementary Fig. 5b).

### Novelty of trait-associated landrace haplotypes

The ultimate criterion for assessing the usefulness of favorable landrace haplotypes for germplasm improvement is their frequency in breeding material. If favorable haplotypes are already present in high frequency in the genetic material to be improved, they are of no additional value. We assessed the frequencies of the identified trait-associated focus haplotypes in a panel of 65 breeding lines based on genotypic data. When tracking an ancestral haplotype potentially shared between landrace and breeding material, recombination might have broken up the respective haplotype, but the trait-associated causal mutation might still be present. Small genetic window sizes (mean = 0.036 cM), low values of historical recombination events (mean = 1.6) and high levels of haplotype similarity (mean = 0.29) found in the panel of breeding lines pointed to a low probability of haplotypes being broken up by recombination.

Frequency distributions of favorable haplotypes in the 65 breeding lines for early development traits (EV_V4, EV_V6, PH_V4 and PH_V6) are given in Fig. 4. As the haplotypes identified for each of the four single traits (Table 1) were partly from similar genomic regions, we only considered 53 favorable haplotypes with a minimum distance of 1 Mb and/or *r*^*2*^ <0.8. The frequency of favorable haplotypes (mean = 0.20) was significantly increased (*P* < 0.01) compared to randomly drawn haplotypes (mean = 0.16). Six favorable focus haplotypes (11%) were absent in the set of breeding lines representing potential novel variation for elite germplasm improvement. The mean frequency of 80 unfavorable haplotypes associated with early plant development did not differ significantly (*P* > 0.30) from the frequency of random haplotypes. A substantial proportion of the unfavorable haplotypes (27.5%) were common in the breeding lines (Fig. 4), suggesting that a targeted substitution with favorable haplotypes could lead to further germplasm improvement.

**Fig. 4:**
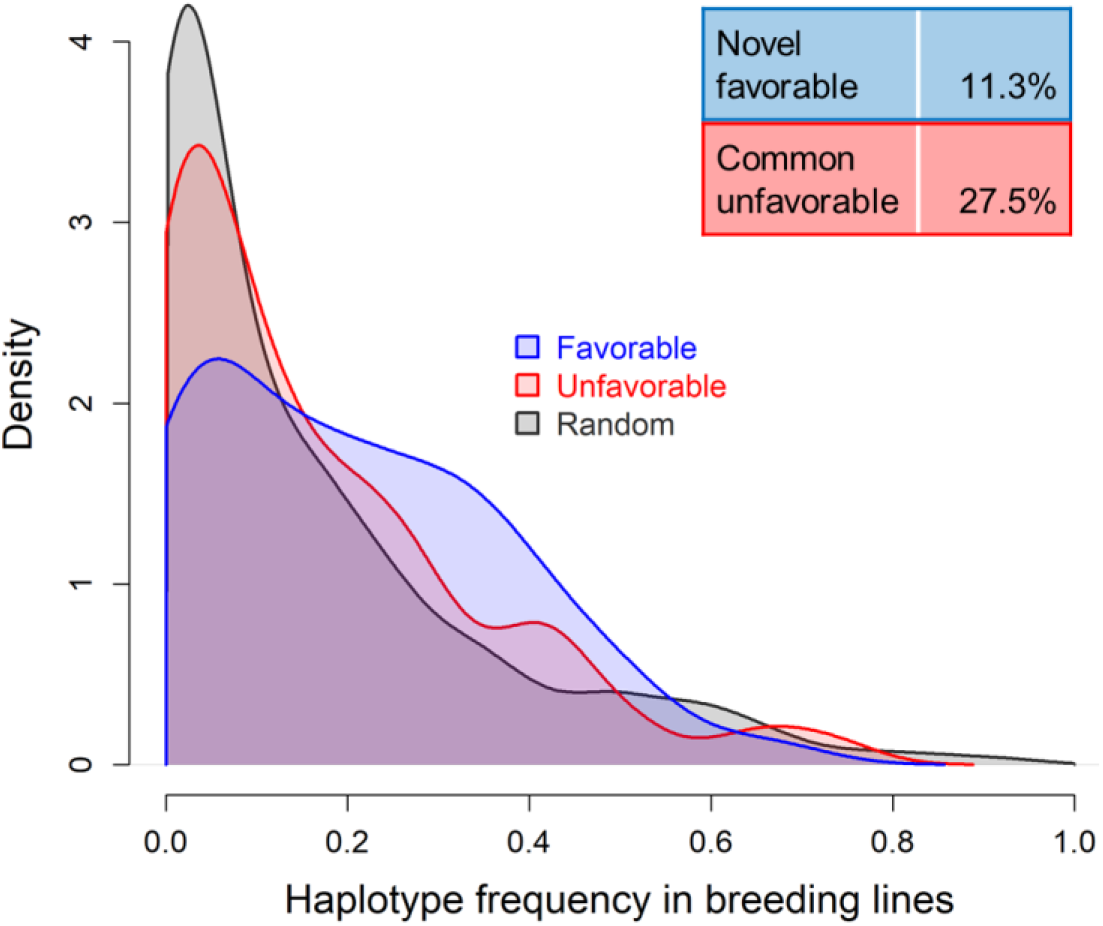
Frequencies of favorable and unfavorable landrace haplotypes in breeding lines. Density estimation for favorable (n = 53, blue), unfavorable (n = 80, red) and random (n = 500, gray) haplotypes in 65 breeding lines. Haplotypes significantly associated with four early plant development traits (EV_V4, EV_V6, PH_V4 and/or PH_V6) in landrace derived DH libraries exhibiting a distance > 1 Mb and/or *r*^*2*^ < 0.8 were considered. Six favorable haplotypes (11.3%) were absent in the breeding lines. 22 unfavorable haplotypes (27.5%) were common in the panel of breeding lines, i.e. having a frequency larger than the upper quartile (>0.231) of random haplotypes.

### Linking novel haplotype variation to phenotypes

To evaluate the potential of individual focus haplotypes to improve elite germplasm, we compared the phenotypic performance of landrace derived DH lines carrying focus haplotypes with a subset of breeding lines (n = 14) tested in six locations in 2017. Exemplarily, we report the results for two genomic regions on chromosomes 3 and 9, found to affect PH_V6 in the GWAS analysis (Fig. 5). On chromosome 3, the focus haplotype (Haplotype A in Fig. 3a and Supplementary Fig. 4) was localized in a 10 SNP window which explained 4.8% of the genetic variation for PH_V6 and comprised eight additional haplotypes in the DH lines. The focus haplotype had a frequency of 4.1% in the DH lines, outperformed six of the eight alternative haplotypes significantly and was absent in the panel of breeding lines. 93.8% of the 65 breeding lines carried one of the six haplotypes with significant negative effects relative to the focus haplotype (on average 0.61 standard deviations) in almost all environments. The remaining breeding lines (6.2%) carried a haplotype absent in the landrace panel and thus without effect estimate. Averaged across environments, DH lines carrying the focus haplotype showed an increase of 6.06 cm over breeding lines, but the difference was not significant (*P* > 0.056; Fig. 5a). When looking at individual environments however, significant differences (*P* < 0.044) were observed for locations OLI, EIN and ROG (Supplementary Fig. 6a), which showed the lowest temperatures in the field^28^ suggesting that the relative advantage of the identified haplotype might be temperature dependent.

**Fig. 5:**
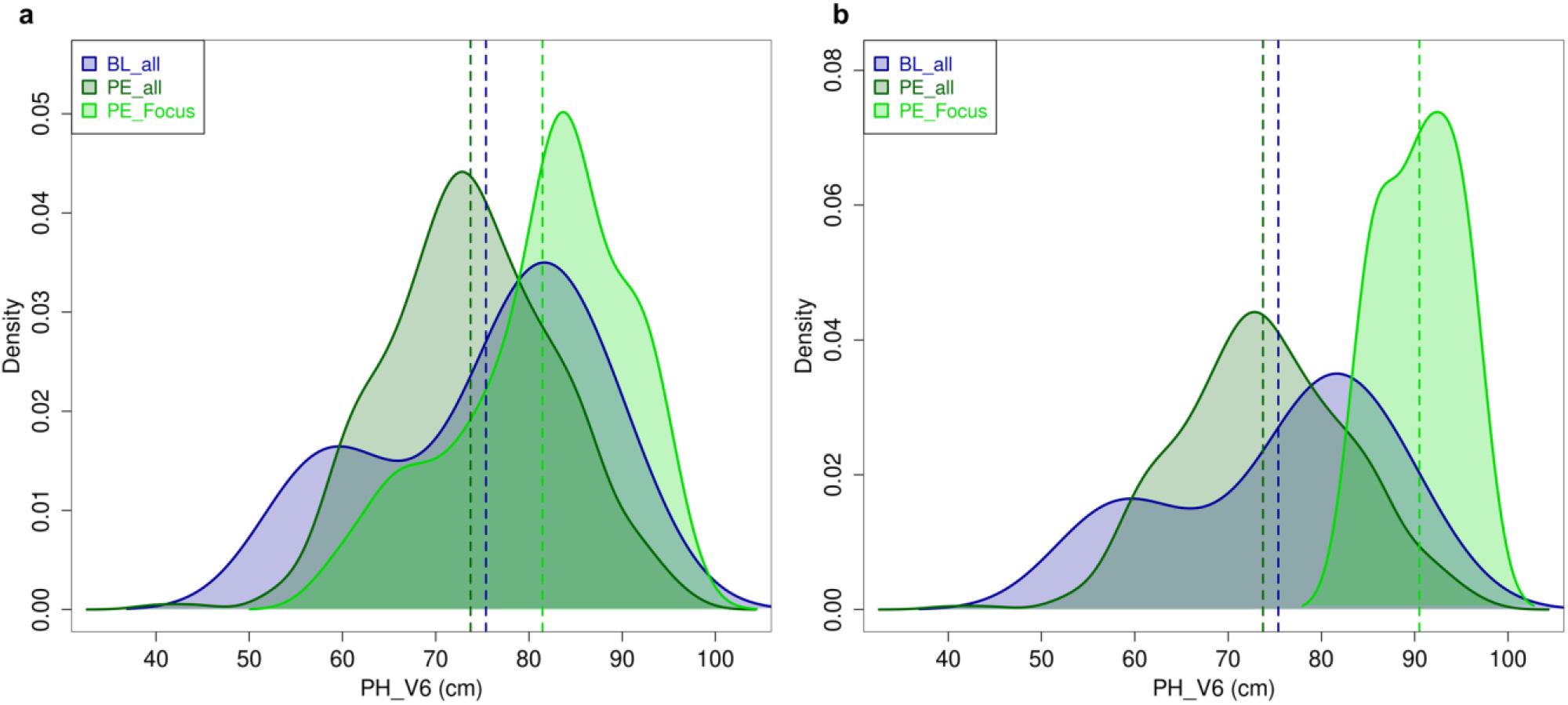
Effect of favorable haplotypes not present in breeding lines on early plant development. Estimated densities of phenotypic values (BLUEs across locations in 2017) for PH_V6 for 14 breeding lines (BL_all), 402 DH lines of landrace PE (PE_all) as well as for DH lines of PE carrying **(a)** a focus haplotype on chromosome 3 (haplotype A in Fig. 3a; PE_Focus, 38 lines) and **(b)** a focus haplotype on chromosome 9 (haplotype D in Fig. 3a; PE_Focus; based on 3 data points only). Vertical lines indicate the mean of each group. The difference in means between BL_all and PE_all was not significant (*P* > 0.514; permutation test).

On chromosome 9 in a genomic region of about 3 Mb, three independent focus haplotypes affected PH_V6 significantly (two favorably, one unfavorably). One of the three focus haplotypes (Haplotype D in Fig. 3a and Supplementary Fig. 4) increased PH_V6 compared to the six alternative haplotypes in the respective window. The genetic variance explained by the haplotypes in this window was small (1.7%) most likely due to the low frequency (0.4%) of the focus haplotype in the DH lines. The focus haplotype was absent in the panel of 65 breeding lines. Instead 95.4% of the breeding lines carried one of the six inferior haplotypes, while 4.6% carried haplotypes not present in the landrace panel. DH lines carrying the focus haplotype showed a significant increase of 15.1 cm compared to the breeding lines (*P* < 0.009). Similar as for the haplotype on chromosome 3, the difference was most pronounced in environments showing low temperature during early plant development (Supplementary Fig. 6b).

We also assessed genomic regions in more detail where the focus haplotype was unfavorable, like for example the window comprising the *tb1* locus which explained 13.1% of the genetic variance for TILL in the landrace panel. DH lines carrying the unfavorable focus haplotype showed a significant increase of 1.51 scores compared to the 14 phenotyped breeding lines not carrying the haplotype (Supplementary Fig. 7a; *P* < 0.0001). Here, the focus haplotype was carried by only two of the 65 breeding lines, but for other genomic regions associated with TILL frequencies were higher, e.g. 15.5% for a region on chromosome 5 explaining 6.6% of the genetic variance in the DH lines. In this case, DH lines carrying the focus haplotype showed a significant increase of 1.69 scores compared to 13 breeding lines not carrying the haplotype (Supplementary Fig. 7b; *P* < 0.0004). For a genomic region on chromosome 1 associated with EV_V4 (Supplementary Fig. 8), more than half of the 65 breeding lines carried the unfavorable focus haplotype, including six of the 14 phenotyped lines. The window in which the focus haplotype was located comprised four additional haplotypes and accounted for 5.1% of the genetic variance in the DH lines. We tested the effect of the focus haplotype in the 14 breeding lines and found a significant difference of 0.875 scores between lines with and without the focus haplotype (*P* < 0.039, Supplementary Fig. 8), indicating that a targeted substitution of the focus haplotype with one of the alternative haplotypes could lead to germplasm improvement.

Introducing novel alleles into elite germplasm for a target trait comes at the risk of undesired effects on other traits due to pleiotropy or linkage. We tested the identified focus haplotypes for each of the early plant development traits in bivariate models for significant effects on other traits (PH_final, FF, MF, LO and TILL,). Of the 53 favorable haplotypes referred to in Fig. 4, 20 had a significant effect on at least one of the five other traits. Thereof, only three haplotypes increased LO or TILL, whereas four haplotypes slightly decreased LO or TILL. Fourteen haplotypes increased PH_final and/or led to earlier flowering whereas one haplotype slightly delayed FF. For some of those haplotypes the effect on traits other than early plant development was substantial (e.g. haplotype “J” in Supplementary Fig. 9a increasing LO). An enrichment of such haplotypes in the breeding germplasm is therefore not advisable. In contrast, haplotypes which explained more of the genetic variance for early plant development than for other traits (e.g. haplotypes “E” or “G” in Supplementary Fig. 9a) still can be used for improving germplasm for early plant development resulting in only slightly altered flowering time and/or PH_final. Of the 80 focus haplotypes unfavorable for early plant development (Fig. 4), 48 were significant for at least one other trait. Thereof, 14 haplotypes decreased TILL, while 40 decreased PH_final and/or delayed flowering. However, most of them had only moderate effects on these traits (Supplementary Fig. 9b). Therefore, in many cases selection against those haplotypes can still be recommended.

## Discussion

The importance of genetic variation for selection and genetic improvement of crops is undisputed. Genetic resources of domesticated species, such as landraces, are a valuable source of diversity for broadening the genetic base of elite germplasm^1^. However, efficient strategies for utilizing this native diversity for the improvement of quantitative traits are lacking. Here, we developed a strategy to discover novel variation for quantitative traits in maize landraces (Supplementary Fig. 1). The combination of comprehensive molecular inventories and meaningful phenotypes collected in landrace derived DH libraries in multi-environment trials allowed detection of novel variation for quantitative traits exhibiting limited genetic variation in elite material. Even though the DH libraries were derived from only three pre-selected populations, 26% of landrace haplotypes were absent in the panel of breeding lines representing the allelic diversity of multiple diverse source populations^27^. While most of these haplotypes can be expected to be neutral^32^ or disadvantageous, some might represent useful novel variation.

Landraces represent self-contained populations adapted to their geographical origin^33^. By focusing on diversity within rather than across landraces, confounding effects of strong adaptive alleles are avoided. Consequently, individual trait-associated haplotypes are expected to have moderate to small effects only. Our results meet these expectations. The majority of haplotype-trait associations detected in the DH libraries explained less than 5% of the genetic variance for all traits under study including flowering time. However, as shown for the haplotype affecting PH_V6 on chromosome 9 (Fig. 5b), the genetic variance explained in GWAS is not only a function of effect size but also of haplotype frequency. As DH and breeding lines were sampled from the same germplasm group (European flint maize), haplotype frequencies were positively correlated between the two panels (Fig. 1b). This exemplifies one of the key challenges when searching for novel variation for quantitative traits, as haplotypes absent in the breeding material tend to have low frequencies also in landraces with shared historical ancestry. Focusing on a set of landraces pre-selected for variation in target traits increases the chances that they harbor alleles at frequencies large enough to be detected in GWAS. The success of this strategy was reflected in the high number of significant haplotype-trait associations found for target traits early vigor and early plant height.

The large sample of landrace derived DH lines employed in this study enabled mapping of haplotypes with only moderate effect size and/or comparably low frequency, but as is known for GWAS studies, some of these significant trait associations might be spurious^34^. Here, the sequential determination of significance (Supplementary Fig. 3) should have minimized the proportion of false positives^35^. In addition, the haplotype-based approach enabled tracking of ancestral alleles between landrace derived and breeding material and the phenotypic comparison between the two groups supported the usefulness for germplasm improvement. Nevertheless, the construction of haplotypes for identification of novel variation in landraces warrants further research. Different methods for haplotype construction exist generating population-specific haplotype blocks based on LD^36,37^ or linkage^38^. Here, we used fixed window sizes, as it is advantageous in comparing haplotype frequencies across datasets varying in their extent of LD. The choice of window size depends on the available marker density and affects the number of haplotypes per window as well as the risk of haplotypes being broken up by recombination. Thus, defining the haplotype inventories of landraces and comparing them to elite germplasm is not trivial. Comprehensive sampling of individuals or lines from a limited number of landraces mitigates difficulties in haplotype construction and at the same time warrants sufficient statistical power and mapping resolution in GWAS through absence of pronounced population structure, rapid decay of LD, and consistency of linkage phases^26^. Here, we put this strategy into practice and showed its potential in identifying novel favorable alleles for improving quantitative traits.

A subset of breeding lines was evaluated together with the DH libraries. For early development traits, overall performance did not differ significantly between the two groups, but DH lines carrying specific focus haplotypes not present in breeding lines outperformed the set of breeding lines significantly in environments favoring trait differentiation. This is a first step in identifying novel haplotypes for germplasm improvement but the final proof of concept will have to come from crosses of landrace derived material with elite material. As landraces represent open-pollinated populations, background dependency of the identified trait-associated haplotypes should not be as pronounced as in mapping populations tracing back to few genetic founders such as multi- or biparental crosses. In our study, the vast majority of trait-associated haplotypes occurring in landraces PE and KE had equal effect signs across landraces and environments supporting this hypothesis. In addition, for cases where it was possible to contrast different haplotypes in the breeding lines (Supplementary Fig. 8), the effect of the focus haplotype in the breeding lines was consistent with the effect in the DH lines. If the selected landraces and the target germplasm to be improved share historical ancestry, we expect only minor genetic background effects when introducing novel variation from landraces into elite material.

After identification of trait associations, fine mapping of the respective genomic regions and functional characterization of candidate genes is a logical next step. With a limited number of annotated genes per trait-associated genomic region, high mapping resolution was obtained in this study. The envisaged functional validation of relevant haplotypes opens many options for utilization: targeted allele mining from genetic resources, unlocking diversity trapped in disadvantageous or incompatible haplotypes, broadening the genetic diversity at relevant loci in elite germplasm and improvement of unfavorable haplotypes through gene editing^39^. In addition to targeted haplotype management, genome-wide approaches will also profit from functional knowledge. Pre-breeding programs^2^ might be accelerated through the use of genome-based prediction^40,41^. It has been shown that prediction accuracy is increased if known trait-associations are included as fixed effects in prediction models^42^. As our results indicate high stability of haplotype effects across environments and genetic background as well as limited haplotype-induced correlations between traits the prospects of germplasm improvement through the use of landrace derived material are promising.

By successfully linking molecular inventories of landraces to meaningful phenotypes and identifying novel favorable variation for quantitative traits of agronomic importance, the results of this study represent a first step towards the long-term goal of accessing native biodiversity in an informed and targeted way. The strategy proposed in this study and demonstrated experimentally with the European flint germplasm can be extended to other maize germplasm groups and even to other allogamous crop species. The key to an efficient use of genetic resources is to understand how genomic information of gene bank accessions can be translated into plant performance^43^. We envision a future where haplotypes characterized for their genomic structure, allele content and functional relevance can be freely moved between populations. Our goal is to create plants with novel combinations of alleles that will lead to varieties with novel combinations of traits, thus securing sustainable crop production in a changing world.

## Methods

### Plant material

We generated more than 1,000 doubled-haploid (DH) lines derived from three European maize landraces: Kemater Landmais Gelb (KE), Lalin (LL) and Petkuser Ferdinand Rot (PE)^28^. The landraces were pre-selected for phenotypic variation in cold-related traits assessed in field trials and population genetic analyses described by Mayer et al.^26^. The set of breeding lines used in this study was based on a broad panel of 68 flint lines described by Unterseer et al.^27^. The initial dataset included two US sweetcorn lines, IL14H and P39, which we excluded in our analyses. The remaining 66 lines, released between ~1950 and 2010, were selected to represent the genetic diversity of the European flint elite breeding germplasm. The panel also includes prominent founder lines like EP1, F2, F7 and DK105^44^.

### Genotypic data

In total, 1,015 landrace derived DH lines were genotyped with the 600k Affymetrix® Axiom® Maize Array^45^. After stringent quality filtering^28^, 941 lines (KE = 501, LL = 31, PE = 409), and 501,124 markers mapped to B73 AGPv4^46^ remained for genetic analyses. Remaining heterozygous calls were set to missing and all missing values were imputed separately for each landrace using Beagle version 5.0^47^ with default settings. From the set of 66 breeding lines, 64 lines were genotyped with the same 600k array^27^, whereas for two lines (EZ5 and F64) overlapping SNP positions (85%) were extracted from the HapMap data^48^ which is based on whole genome sequences. For making the 600k genotyping data comparable to the HapMap data, all alleles were coded according to the B73 AGPv4^46^ forward strand. The breeding line data was filtered for the 501,124 high quality markers of the set of DH lines. Applying the same quality filter criteria as for the DH panel (heterozygous calls < 5%; callrate > 90%, except for EZ5 and F64 with callrate >84%), one breeding line (FV66) was removed due to an increased number of heterozygous calls. For the remaining 65 lines heterozygous calls were again set to missing and missing values imputed using Beagle version 5.0^47^ with default settings. For the combined set of landrace derived DH lines and breeding lines, principal coordinate analysis^49^ (PCoA) was conducted based on modified Rogers’ distances^50^ (MRD). Pairwise *r*^*2*^ ^51^ between SNPs within 1 Mb distance was calculated for the DH libraries (within and across the three landraces) and the panel of breeding lines, respectively. Average LD decay distance (*r*^*2*^ < 0.2) was estimated using non-linear regression^52^. If not denoted otherwise, analyses were done using R version 3.6.0^53^.

### Phenotypic data

In total, 958 DH lines were phenotyped for various traits over two years in up to eleven environments, as described by Hölker et al.^28^. A subset of nine traits was analyzed in this study (Supplementary Table 2), related to early plant development, maturity as well as agronomic characteristics. After stringent quality filtering^28^, phenotypic data of 899 DH lines (KE = 471, LL = 26, PE = 402) remained for further analyses. Additionally, 14 checks, comprising representative lines of the European flint breeding pool and included in the panel of 65 genotyped breeding lines, were phenotyped in six locations in 2017^28^. Best linear unbiased estimates (BLUEs) for each DH line and check were calculated across environments using a mixed linear model as described by Hölker et al.^28^. Analogously, BLUEs were calculated within each environment using the same model without environment related model terms.

### Haplotype construction

For both, the landrace derived DH lines as well as the breeding lines, haplotypes were defined as a given nucleotide sequence within non-overlapping windows of 10 SNPs (Supplementary Fig. 3a). For the 600k chip, the density of SNPs along the chromosomes follows the average recombination rate^45^. Therefore, using a fixed number of SNPs per window leads to similar window sizes as defined based on genetic map units. The median physical window size was 17.2 kb (mean = 40.7 kb), corresponding to 0.008 cM (mean = 0.036 cM) according to a genetic map generated from a F2 mapping population of a cross of EP1×PH207^44^. Within each window, haplotypes were coded as presence/absence markers, yielding genotype scores 0 and 2 for DH and breeding lines. To evaluate the potential of novel variation in landraces, we compared haplotype frequencies between the landrace derived DH lines and the panel of 65 breeding lines.

### Identification of trait-associated haplotypes

For GWAS in the DH lines, haplotypes which were present less than three times in the panel of 899 phenotyped DH lines were excluded from the analysis. For haplotypes with *r*^*2*^ = 1, only one was retained, resulting in 153,730 haplotypes used for GWAS (Supplementary Fig. 3a), with on average 5.73 haplotypes per window. The identification of trait-associated haplotypes was conducted in two steps following Millet et al.^35^, (i) identification of candidate haplotypes in GWAS (Supplementary Fig. 3b) and (ii) backward elimination in a multi-locus multi-environment model (Supplementary Fig. 3c). GWAS were conducted for single environments as well as across environments using the corresponding environment-specific and across-environment BLUEs as response variable in the model, respectively. A univariate linear mixed model, implemented in GEMMA version 0.98.1^54^, was used:

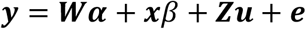

where *y* is the *n*-dimensional vector of phenotypic values (BLUEs), with *n* being the number of lines; *α* is a three-dimensional vector of fixed effects (intercept and landrace effects of KE and LL); *β* is the fixed effect of the tested haplotype; *x* is the vector of corresponding genotype scores coded as 0 and 2; *u* is the *n*-dimensional vector of random genotypic effects, with 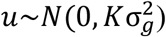; and *e* is the *n*-dimensional vector of random residual effects, with *e*~*N*(0, *I*_*n*_σ^2^). *K* denotes the (*n* × *n*) genomic relationship matrix based on SNP markers according to Astle and Balding^55^ and *I_n_* the (*n* × *n*) identity matrix. Matrices *W* (*n* × *3*) and Z (*n* × *n*) assign phenotypic values to fixed and random effects, respectively. Significance of haplotype-trait associations was assessed for each single-environment as well as for the across-environment GWAS based on the likelihood ratio test, as implemented in GEMMA, using a 15% false discovery rate^56^ (FDR). Haplotypes with a physical distance of less than 1 Mb and in high LD (*r*^*2*^ ≥ 0.8) were considered to mark the same genomic region. The corresponding trait-associated genomic region was described by the start and end positions of the first and last haplotype fulfilling the defined criteria. To represent genomic regions equally in subsequent analyses, only the most significant haplotype, the focus haplotype, was retained per region in the respective GWAS, resulting in a set of candidate haplotypes.

In the multi-locus, multi-environment (MLME) mixed linear model, we conducted a backward elimination of those candidate haplotypes as suggested by Millet et al.^35^, using the ASReml-R package version 3.0^57^:

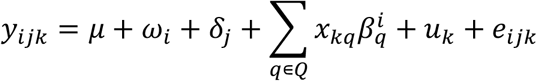

where *y*_*ijk*_ is the phenotypic value (BLUE) of line *k* belonging to landrace *j* tested in environment *i*; *μ* is the common intercept; *ω*_*i*_ is the fixed effect of environment *i*; *δ*_*j*_ is the fixed effect of landrace *j*; *x*_*kq*_ is the genotype score (0 or 2) of line *k* for haplotype *q*; 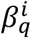 is the fixed effect of haplotype *q* in environment *i* comprising the haplotype main and haplotype by environment interaction effect, i.e. 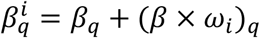; *u*_*k*_ is the random genotypic effect of line *k*, and *e*_*ijk*_ is the random residual error with environment-specific residual error variance. *Q* represents the final set of haplotypes obtained through step-wise backward elimination based on the Wald test for 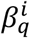 ^58^. At each step, significance of each haplotype was tested when it was the last one entering the model and the least significant haplotype was removed if *P* ≥ 0.01. The proportion of genetic variance explained by the set of trait-associated haplotypes was estimated by calculating the reduction in 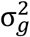 between models including and excluding the haplotype effects, following Millet et al.^35^. For evaluating effect stability across landraces for the final set of haplotypes *Q*, we estimated landrace-specific haplotype effects for each environment using the same MLME model but changing the term 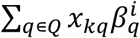 to 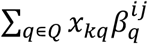, with 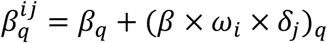.

### Favorable and unfavorable haplotypes and their effect stability across environments

The number of environments in which a haplotype was significant was estimated by generating 95% confidence intervals (CI = effect estimate ± 1.96 × standard error) based on the MLME model, following Millet et al.^35^. A CI not including 0 indicated significance of the haplotype in a given environment. Haplotypes with constant effect sign across significant environments, were classified as “favorable” or “unfavorable”. For EV_V4, EV_V6, PH_V4 and PH_V6 positive (negative) effects were defined as favorable (unfavorable). For LO and TILL negative (positive) effects were defined as favorable (unfavorable). No classification was made for PH_final, FF and MF, as breeding goals vary for these traits. Haplotypes with changing sign of significant effects in different environments were classified as “interacting”.

### Haplotypes associated with multiple traits

For pairwise combinations of early plant development traits with other traits, we tested if haplotypes identified for early plant development also had an effect on the respective other trait using a bivariate model, similar to Stich et al.^59^:

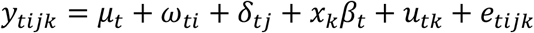

where, *y*_*tijk*_ is the phenotypic value (BLUE) for trait *t* of line *k* belonging to landrace *j* tested in environment *i*; *μ*_*t*_ is the intercept for trait *t*; *ω*_*ti*_ is the fixed effect of environment *i* for trait *t*; *δ*_*tj*_ is the fixed effect of landrace *j* for trait *t*; *x*_*k*_ is the genotype score (0 or 2) of line *k* for the tested haplotype; *β*_*t*_ is the fixed effect of the haplotype for trait *t*; *u*_*tk*_ is the random genotypic effect of line *k* for trait *t*, with *u*~*N*(0, *G* ⊗ *K*); and *e*_*tijk*_ is the residual with *e*~*N*(0, *E* ⊗ *I*_*n*_). *G* and *E* correspond to the (*t* × *t*) genomic and error variance-covariance matrices among traits, respectively, and ⊗ denotes the Kronecker product. Haplotypes for which the 95% CIs for both *β*_*t*_ did not include 0 were considered significant for both traits. The proportion of genetic variance explained per trait by significant haplotypes was estimated by calculating the respective reduction in *G* between models including and excluding the haplotype.

### Comparison of trait-associated haplotypes between landraces and breeding lines

We assessed frequency distributions of identified trait-associated favorable and unfavorable landrace haplotypes in the panel of 65 breeding lines and compared them with 500 haplotypes randomly drawn out of the set of haplotypes occurring at least three times in the landrace panel. Significance for differences in means between the frequencies of favorable and random haplotypes as well as unfavorable and random haplotypes was tested with the Mann-Whitney test. When tracking potentially shared ancestral haplotypes between populations, the probability of a haplotype being broken up by recombination depends on the haplotype length, the recombination rate in the respective genomic region and the time since the most recent common ancestor potentially carrying that haplotype. To evaluate to which extend recombination might have occurred in the haplotypes constructed in this study, we considered the physical as well as genetic length of each haplotype and calculated haplotype similarity (1 – haplotype heterozygosity^60^) and the minimum number of historical recombination events^61^ within the respective genomic windows.

To evaluate the effect of the selected focus haplotype relative to the alternative haplotypes in a given 10 SNP window, we followed the approach of Bustos-Korts et al.^62^, changing the MLME model described above to:

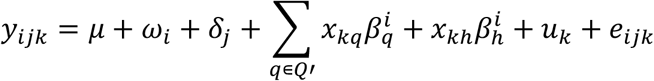

where *Q*′ represents the set of haplotypes *Q* as described above without the respective focus haplotype of the window tested, *x*_*k*ℎ_ is the vector of haplotypes (categorical variable) in the window tested and 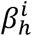 represents the effect of each haplotype in that window relative to the focus haplotype. Similar as above, significance of haplotype effects relative to the focus haplotype was determined by constructing 95% CIs. We further estimated the proportion of genetic variance explained by the given window by calculating the reduction in 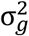 between the null model (without 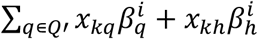) and the model with the 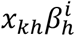 term.

To evaluate to which extent haplotypes with favorable or unfavorable effects in landraces have also favorable or unfavorable effects in elite material, respectively, we compared performance levels between the landrace derived DH lines and the 14 breeding lines used as checks. As phenotypic data for the 14 breeding lines were only available for 2017, only the six environments from 2017 were considered. For some traits differences in means between the landraces were observed^28^, thus comparisons were conducted for each landrace separately. Significance for differences in means between the respective landrace and the 14 checks was tested based on 10,000 permutations (two-sided test). In addition to a comparison of the overall performance level between all lines of the respective landrace and the 14 breeding lines, we compared means between groups of lines carrying a particular haplotype and lines not carrying the haplotype.

## Data availability

Seeds from all genotypes used in the study are available through material transfer agreements. The genotypic data of 941 DH lines and the phenotypic data of 899 DH lines and 14 breeding lines are available at https://doi.org/10.6084/m9.figshare.12137142 (after the manuscript is accepted for publication in peer-reviewed journal). The 600k data of 63 breeding lines can be accessed at https://dx.doi.org/10.6084/m9.figshare.3427040.v1, while for two lines genotypic data based on whole genome sequences were downloaded from http://cbsusrv04.tc.cornell.edu/users/panzea/download.aspx?filegroupid=34.

## Acknowledgements

We are indebted to the technical staff at KWS SAAT SE & Co. KGaA (KWS), Misión Biológica de Galicia, Spanish National Research Council (CSIC), Technical University of Munich, and University of Hohenheim for their skilled support in carrying out the extensive phenotypic evaluation required for this study. We are grateful to Therese Bolduan (KWS) and Tanja Rettig (KWS) for their contribution in planning and conducting field trials. We thank the technical staff at KWS for DNA extraction as well as Hans Rudolf Fries (Technical University of Munich) for processing the genotyping arrays and Stefan Schwertfirm for technical assistance. This study was funded by the Federal Ministry of Education and Research (BMBF, Germany) within the scope of the funding initiative “Plant Breeding Research for the Bioeconomy” (Funding ID: 031B0195, project “MAZE”) and by KWS under Ph.D. fellowships for Manfred Mayer and Armin C. Hölker.

## Author Contributions

CCS and MO, conceived the study; CCS, MO, and AEM acquired funding for the study; MM, ACH, TP, MO, AEM and CCS generated phenotypic and genotypic data; ACH contributed to analyses of phenotypic data; EG contributed to haplotype construction; MM performed analyses and drafted the manuscript; CCS edited the manuscript; all authors read and approved the final manuscript.

## Supplementary Figures

**Supplementary Fig. 1:**
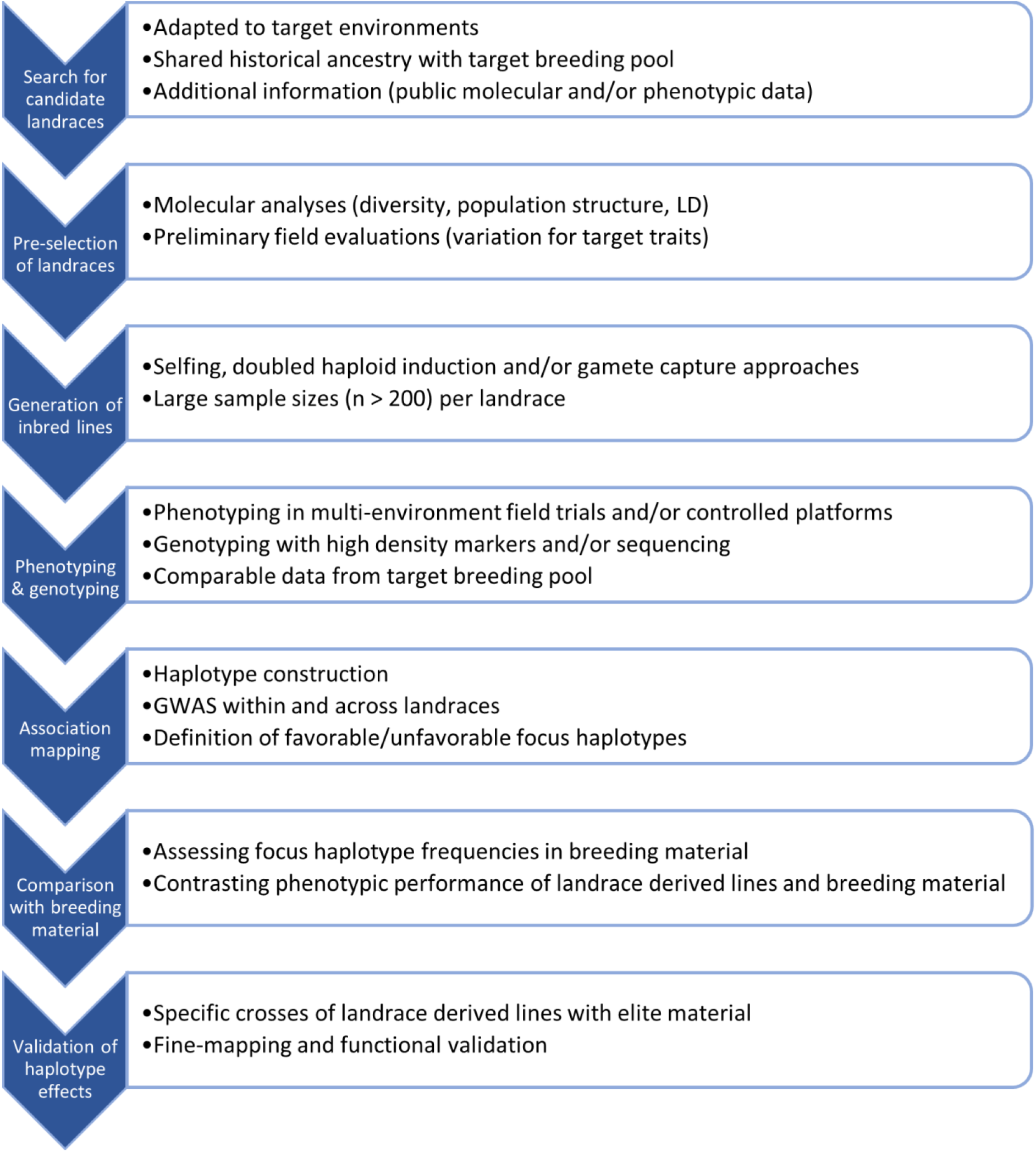
Workflow for making native diversity of landraces accessible for the improvement of elite germplasm.

**Supplementary Fig. 2:**
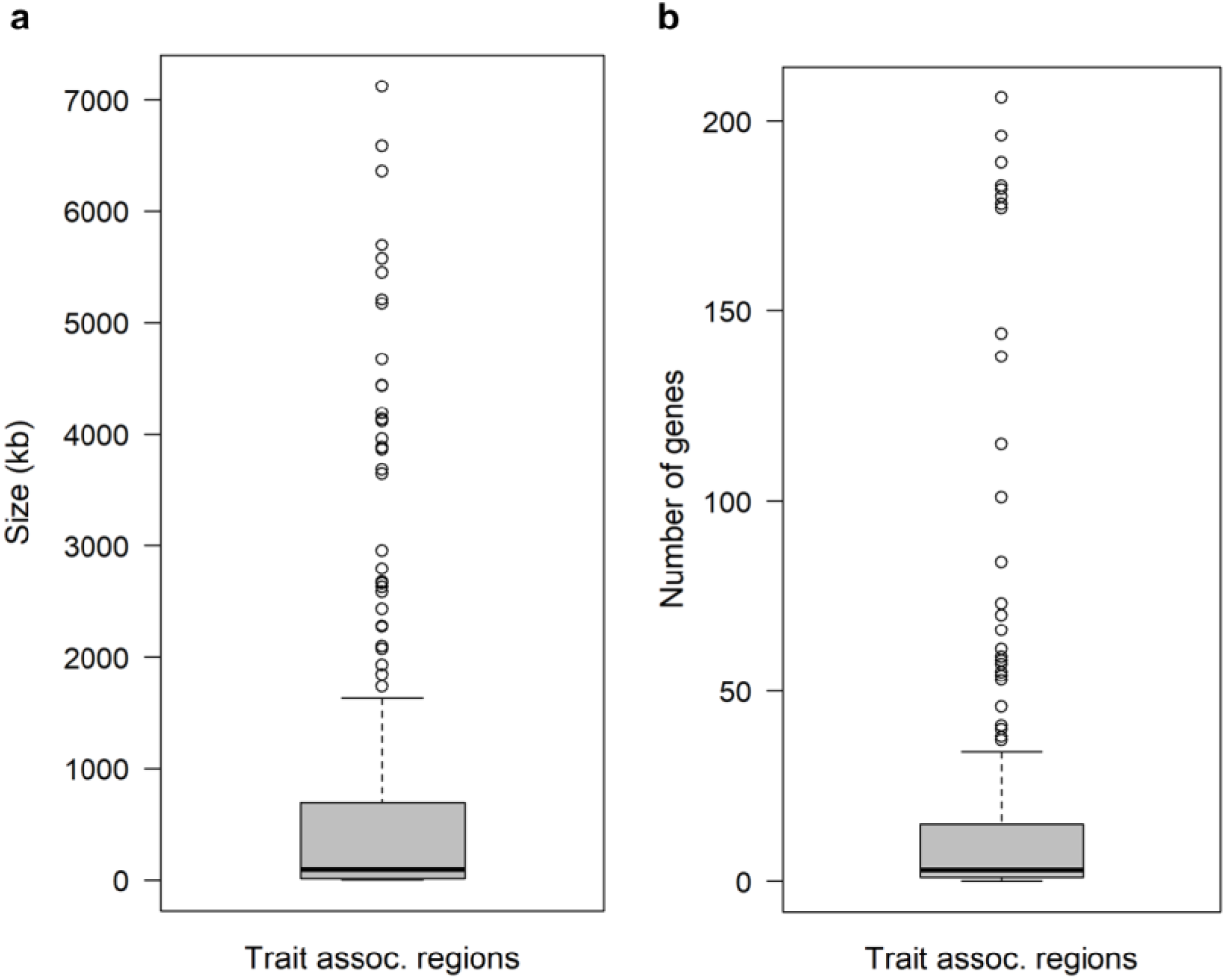
Size (a) and number of annotated genes (b) of/in trait-associated genomic regions. In total 324 genomic regions associated with the traits EV_V4, EV_V6, PH_V4, PH_V6, PH_final, FF, MF, LO or TILL were discovered in 899 DH lines derived from three maize landraces.

**Supplementary Fig. 3:**
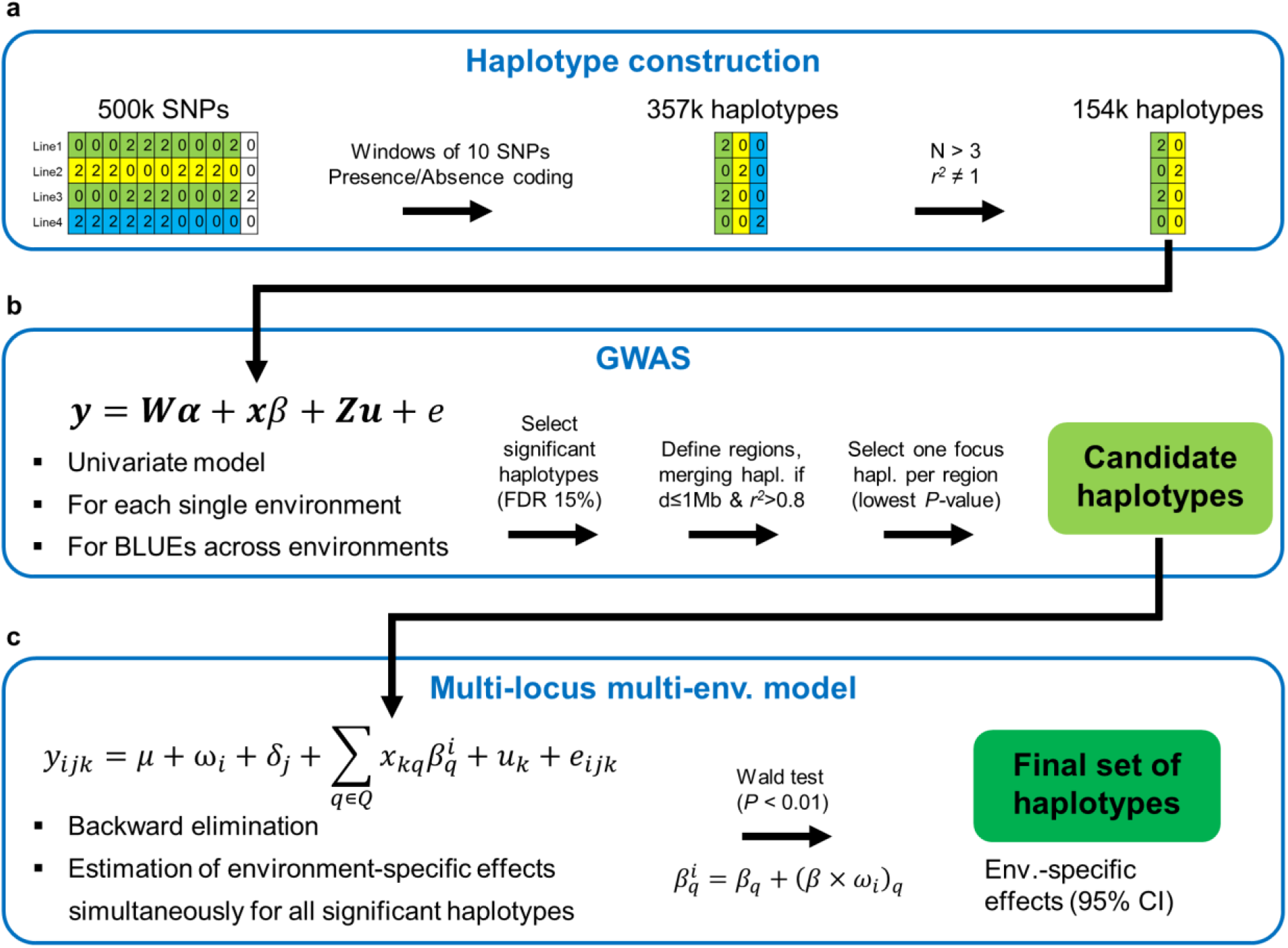
Flowchart of experimental analyses. **(a)** Construction of haplotypes. Haplotype numbers are according to the DH panel. **(b)** GWAS conducted for up to 11 single environments as well as for the across environment BLUEs for the combined set of 899 genotyped DH lines derived from three landraces. **(c)** Multi-locus, multi-environment model for performing backward elimination of candidate haplotypes (Wald test) and estimating environment-specific haplotype effects for the final set Q of focus haplotypes.

**Supplementary Fig. 4:**
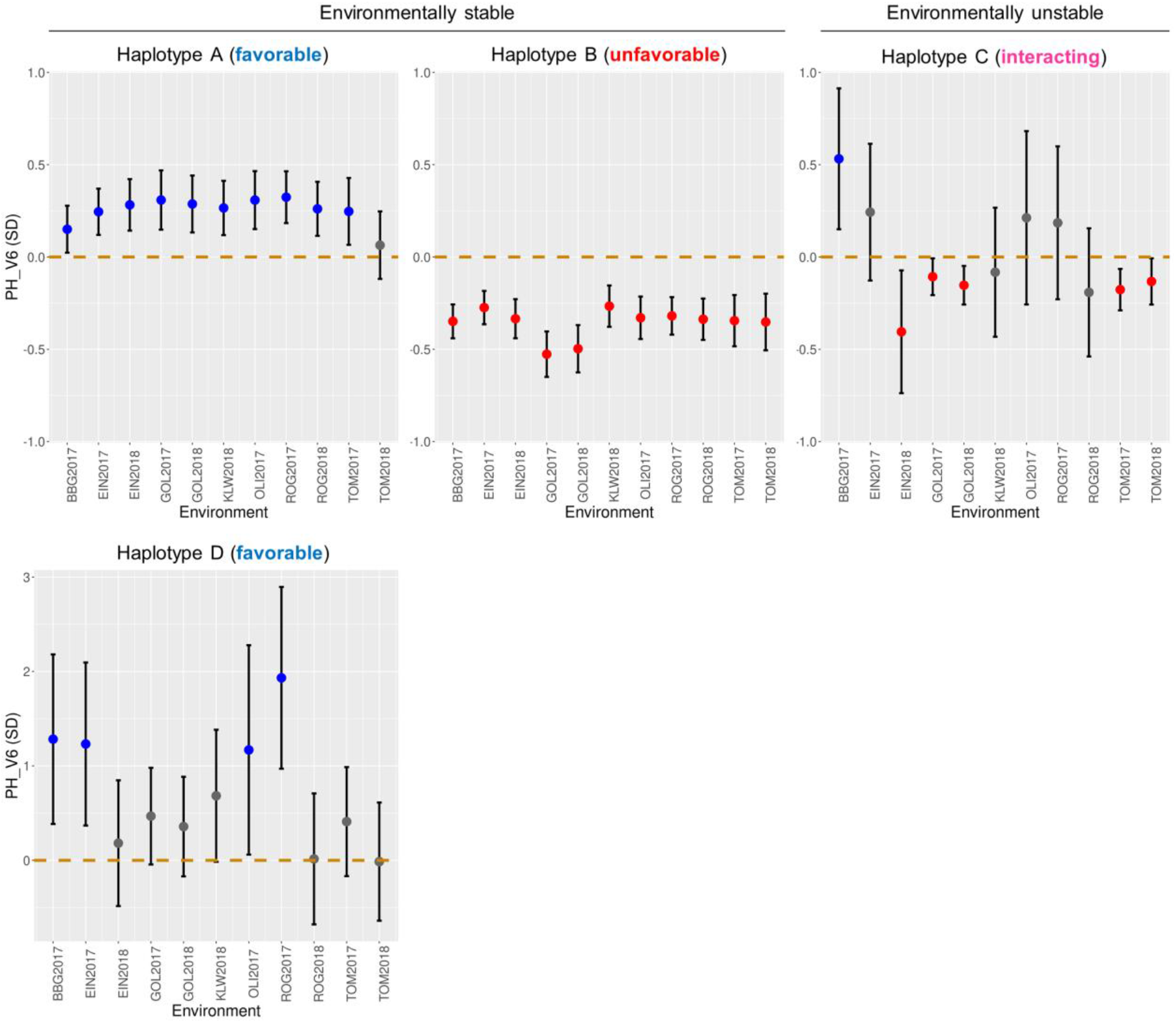
Examples of favorable, unfavorable and interacting haplotypes. Environment-specific effect estimates in units of phenotypic standard deviations with corresponding 95% confidence intervals for four haplotypes associated with PH_V6 (positive, negative and non-significant effects in blue, red and gray, respectively). Haplotype A on chromosome 3 showed significant positive effects in ten out of eleven environments increasing early plant growth, thus representing an environmentally stable favorable haplotype. Haplotype B on chromosome 10 showed negative effects across all eleven environments, decreasing early plant growth thus categorized as environmentally stable unfavorable haplotype. The effect sign of haplotype C on chromosome 9 varied depending on the environment and thus it was categorized as interacting. Haplotype D on chromosome 9 represents another example of an environmentally stable favorable haplotype, showing positive effects in all environments where the association was significant.

**Supplementary Fig. 5:**
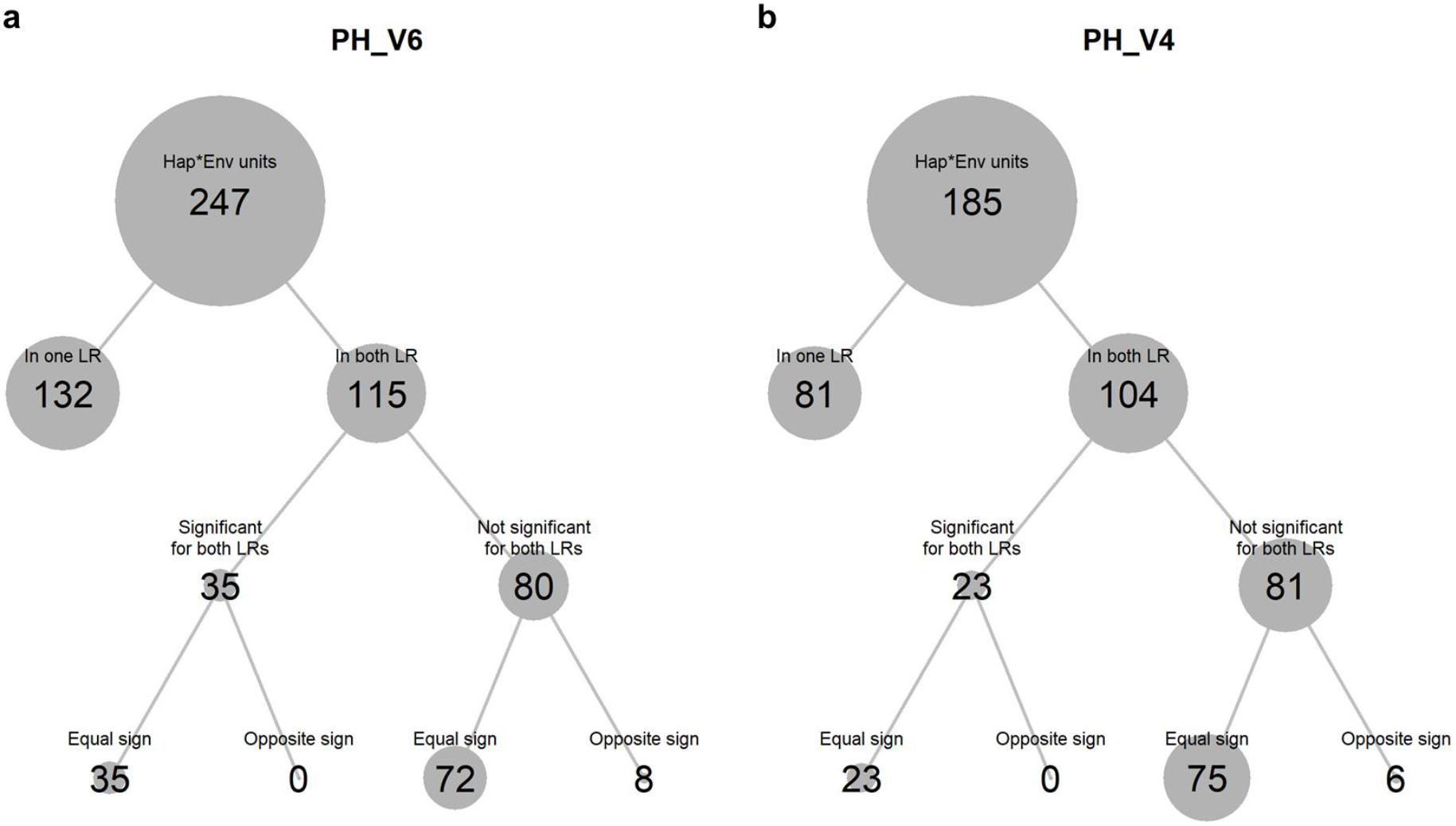
Comparison of haplotype effects between landraces. **(a)** For PH_V6, 46 of the 48 significantly associated haplotypes were present in at least one of the two landraces KE and PE (two occurred only in LL). The sum of haplotype by environment combinations for which the respective haplotypes were significant was 247. Thereof, 132 environment-specific associations resulted from 27 haplotypes only present in either KE or PE, while 115 associations resulted from 19 haplotypes present in both landraces. Thereof, 35 associations (involving 12 haplotypes) were significant for both landraces, whereas 80 associations were significant for one landrace. All 35 associations significant for both landraces had equal effect signs for both landraces. For the 80 associations only significant for one landrace, 72 had equal effect signs for both landraces. **(b)** Analogous to (a) for PH_V4. Here, 36 haplotypes associated to PH_V4 were present in KE and/or PE. Thereof, 21 haplotypes were present in both KE and PE. The 23 environment-specific associations significant for both landraces involved 13 haplotypes.

**Supplementary Fig. 6:**
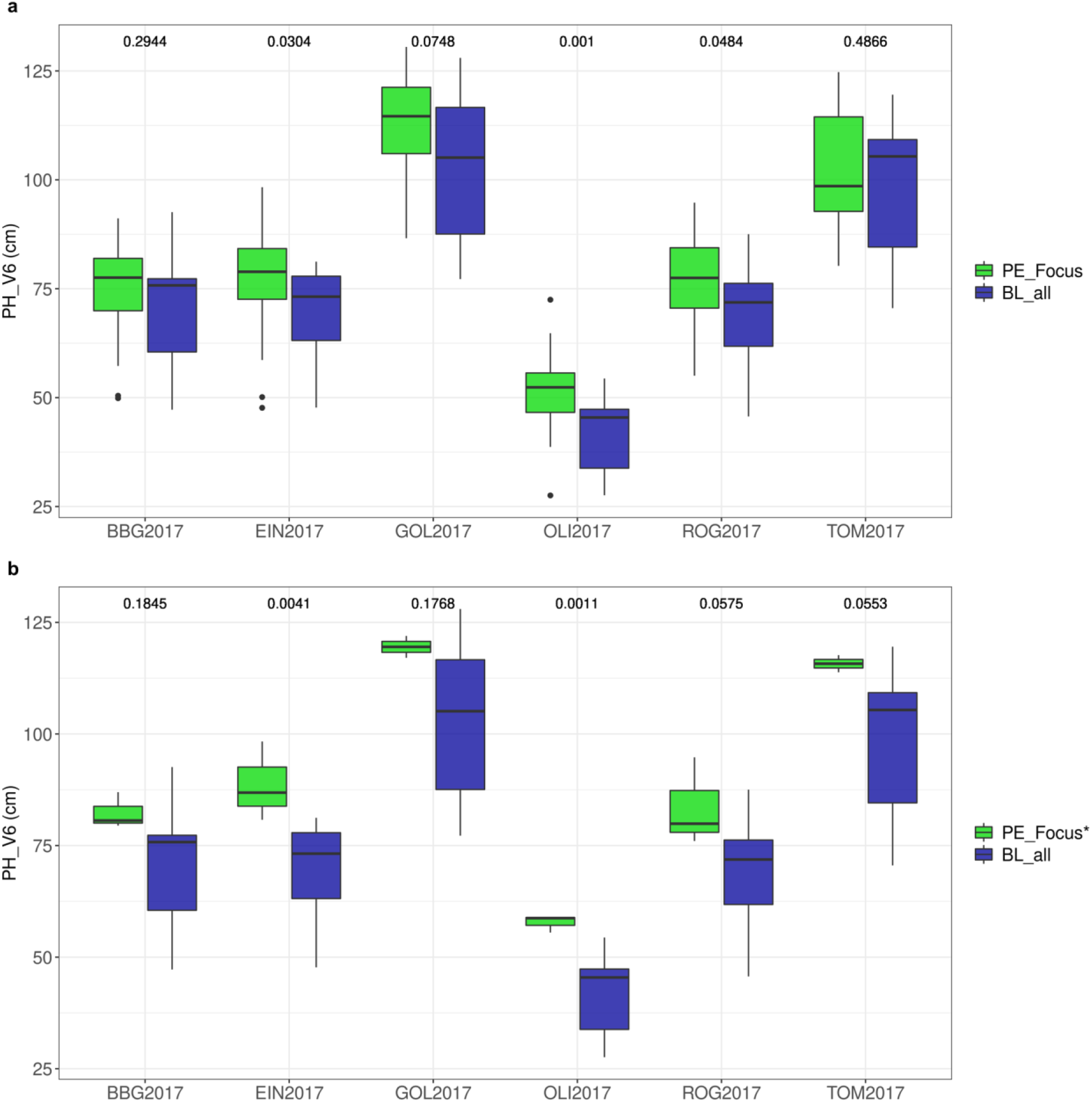
Phenotypic values for PH_V6 in six locations in 2017 for 14 breeding lines (BL_all) and a subset of DH lines derived from the landrace PE (PE_Focus) carrying a focus haplotype. The focus haplotypes in **(a)** and **(b)** refer to haplotype A and D described in Supplementary Fig. 4, respectively. Numbers on the top refer to *P*-values testing the significance of mean differences between BL_all and PE_Focus (permutation test). *based on 3 data points only.

**Supplementary Fig. 7:**
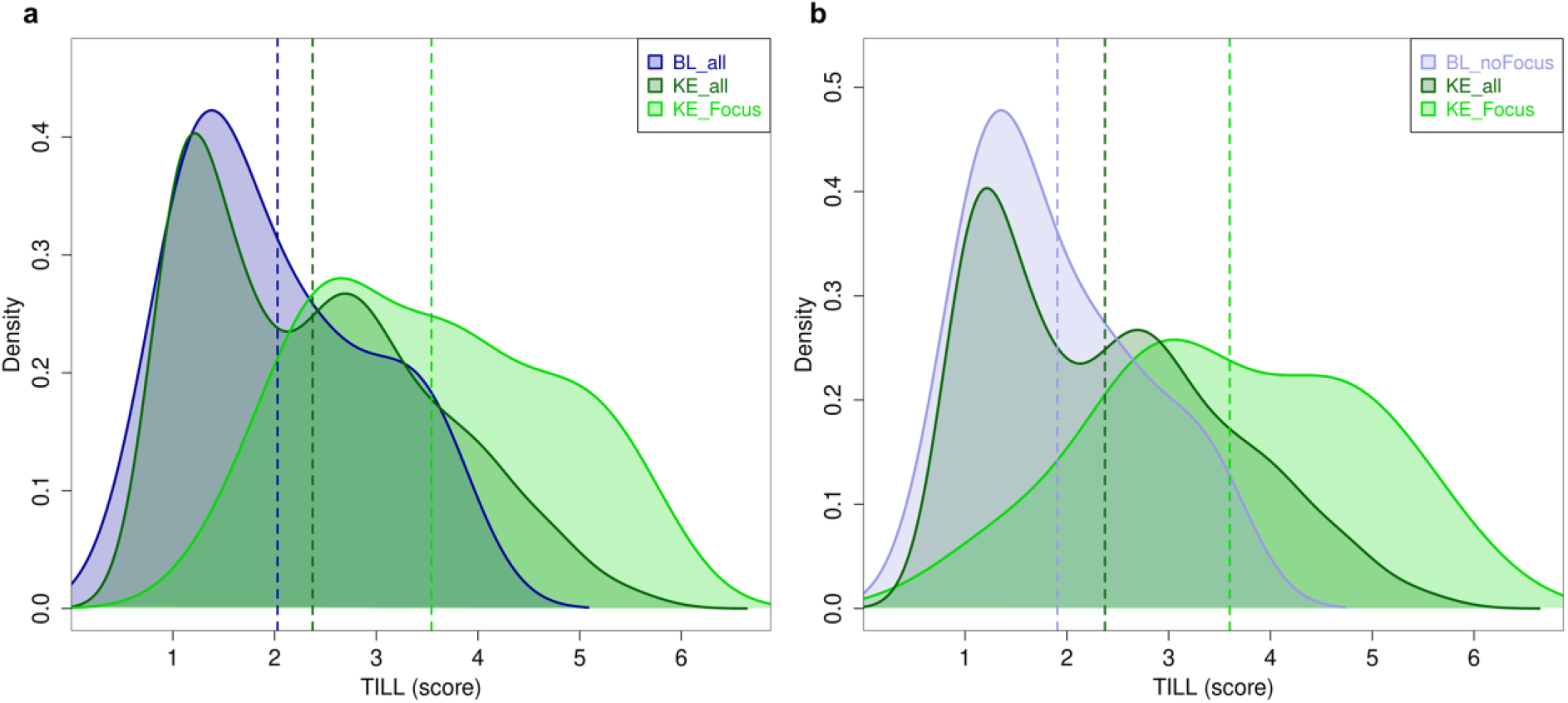
Unfavorable focus haplotypes increasing TILL. Estimated densities of phenotypic values (BLUEs across locations in 2017) for TILL for 14 breeding lines (BL_all), 471 DH lines of landrace KE (KE_all) as well as for DH lines of KE carrying **(a)** a focus haplotype on chromosome 1 (at the *tb1* locus; KE_Focus, 35 lines) and **(b)** a focus haplotype on chromosome 5 (KE_Focus, 16 lines). As one of the 14 breeding lines carried the focus haplotype on chromosome 5, comparisons were made with the remaining 13 lines (BL_noFocus). Vertical lines indicate the mean of each group. The difference in means between BL_all and KE_all was not significant (*P* > 0.277; permutation test).

**Supplementary Fig. 8:**
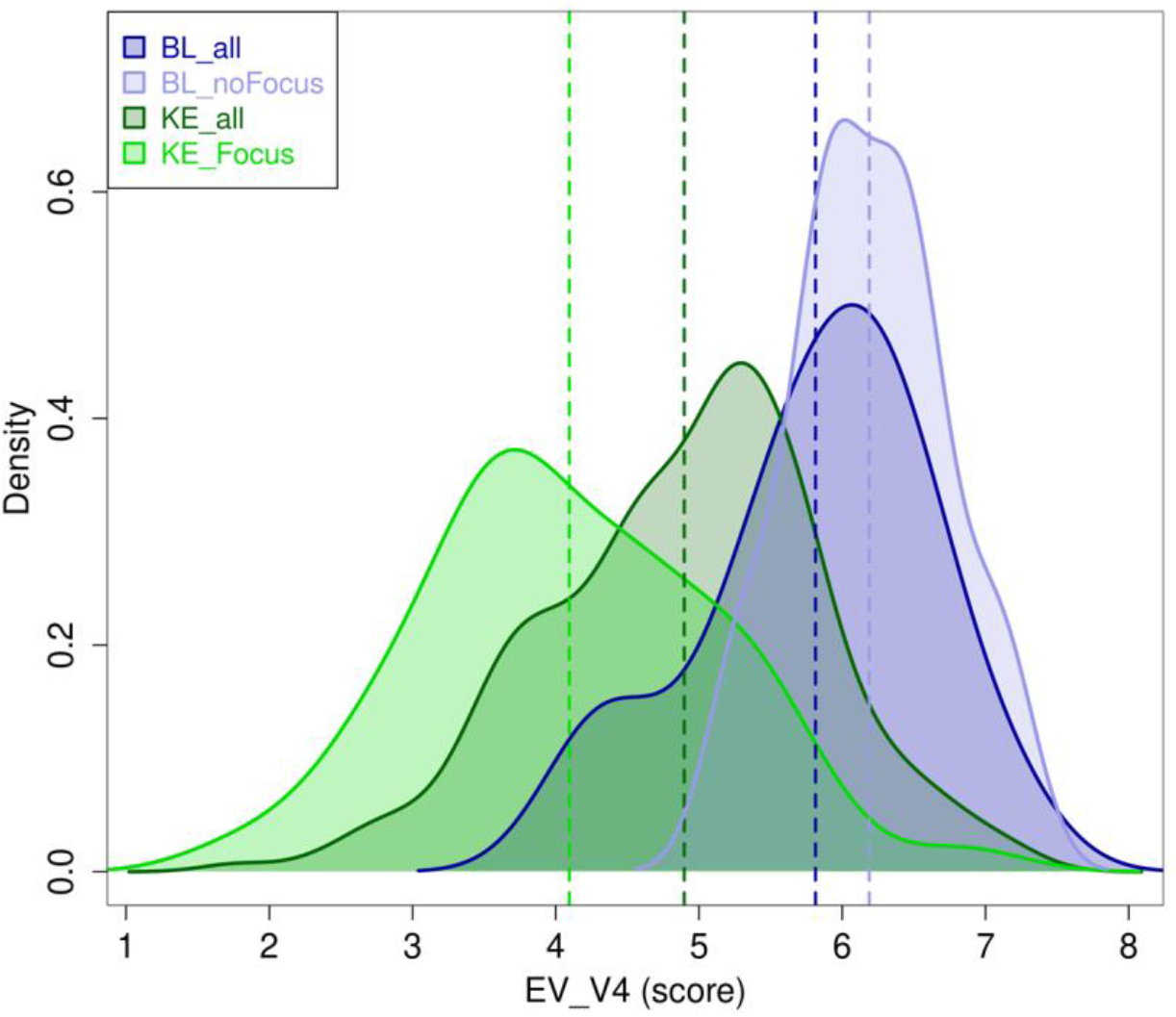
Unfavorable focus haplotype decreasing EV_V4 in landraces and breeding lines. Estimated densities of phenotypic values (BLUEs across locations in 2017) for EV_V4 for 14 breeding lines (BL_all), 471 DH lines of landrace KE (KE_all) as well as for 49 DH lines of KE (KE_Focus) carrying and eight breeding lines (BL_noFocus) not carrying the focus haplotype on chromosome 1. Vertical lines indicate the mean of each group.

**Supplementary Fig. 9:**
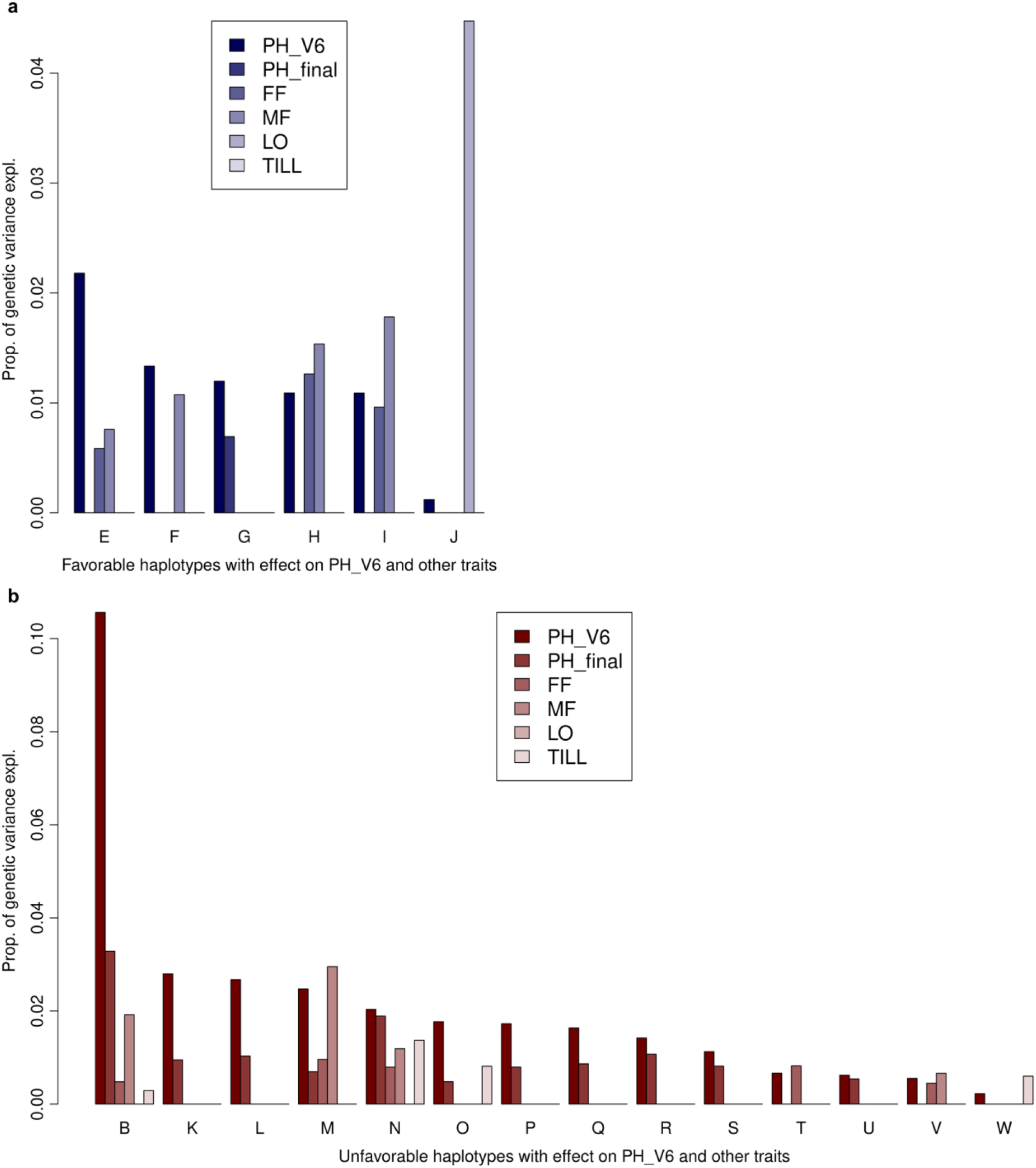
Haplotypes with effects on multiple traits. Proportions of explained genetic variance per trait for each of the six favorable **(a)** and 14 unfavorable **(b)** focus haplotypes associated with PH_V6 with significant effects on PH_final, FF, MF, LO, and/or TILL. Bars are only shown for traits for which the respective haplotype was significant. All of these haplotypes had equal effect signs for PH_V6 and PH_final/LO/TILL and opposite effect signs for PH_V6 and FF/MF, respectively.

## Supplementary Tables

**Supplementary Table 1:** List of identified trait associations. Lists for each trait the position (chromosome, start and end) and size (kb) of each identified trait-associated genomic region as well as the start and end positions of the selected focus haplotype. Further, the range of the corresponding environment-specific haplotype effects (in units of environment-specific standard deviations) together with the respective environments in which the minimum and maximum effects were observed are shown. The table also states for each region the number of environments in which the focus haplotype had a significant effect. Further, the number of annotated genes within each region (including segments 5 kb upstream of genes) according to the B73 AGPv4 reference sequence are shown.

**Supplementary Table 2:** Overview of the phenotypic data analyzed in this study. Lists for each trait, the abbreviation, the way of measurement, the growth stages at which measurements were conducted, the number of lines for which data was available and the number of environments in which the traits were measured.

| Trait | Abbr. | Measurement | Growth Stages | N DHs | N Env. |
| --- | --- | --- | --- | --- | --- |
| Early plant height | PH | Total height in cm, from soil surface to highest tip of upwards stretched leaves, mean of three representative plants per plot | V4, V6 | 899 | 11 |
| Early vigor | EV | Score 1-9, visual appearance of whole plot, 1 = very small plants with discolored leaves, 9 = very vigorous and healthy looking plants | V4, V6 | 899 | 11 |
| Final plant height | PH_final | Total height in cm, from soil surface to lowest tassel branch, mean of three representative plants per plot | R4 | 899 | 11 |
| Female flowering | FF | Days after sowing until 50% of plants show silks | R1 | 899 | 10 |
| Male flowering | MF | Days after sowing until 50% of plants shed pollen | R1 | 899 | 5 |
| Lodging | LO | Score 1-9, 1 = no lodging, 9 = all plants show severe lodging | R6 | 869 | 4 |
| Tillering | TILL | Score 1-9, 1 = no tillers, 9 = many and long tillers | V8-V10 | 899 | 5 |

